# Phospholipid flipping facilitates annexin translocation across membranes

**DOI:** 10.1101/241976

**Authors:** Sarah E Stewart, Avraham Ashkenazi, Athena Williamson, David C Rubinsztein, Kevin Moreau

## Abstract

Annexins are phospholipid binding proteins that somehow translocate from the inner leaflet of the plasma membrane to the outer leaflet. For example, Annexin A2 is known to localise to the outer leaflet of the plasma membrane (cell surface) where it is involved in plasminogen activation leading to fibrinolysis and cell migration, among other functions. Despite having well described extracellular functions, the mechanism of annexin transport from the cytoplasmic inner leaflet to the extracellular outer leaflet of the plasma membrane remains unclear. Here, we show that phospholipid flipping activity is crucial for the transport of annexins A2 and A5 across membranes in cells and in liposomes. We identified TMEM16F (anoctamin-6) as a lipid scramblase required for transport of these annexins to the outer leaflet of the plasma membrane. This work reveals a mechanism for annexin translocation across membranes which depends on plasma membrane phospholipid flipping.

## Results and discussion

As annexin A2 and A5 bind to negatively charged lipid head groups of the inner and outer leaflets of the plasma membrane in a calcium-dependent manner ^1-4^, we hypothesised that lipid remodelling may be involved in their translocation to the cell surface. To assess the role of lipid remodelling in annexin transport across membranes, we studied the effect of the lipid flipping peptide, cinnamycin, in mammalian cells ^5,6^. Cinnamycin is a 19-amino acid lantibiotic that flips phosphatidylethanolamine (PE) and phosphatidylserine (PS) from one leaflet of the lipid bilayer to the other, both in liposomes and cell membranes ^5,6^. We confirmed that cinnamycin flips lipids in HeLa cells, as measured by the amount of PE and PS on the cell surface determined by flow cytometry (Fig. 1a). We next treated HeLa cells with cinnamycin for 30 min in serum-free medium (SFM), washed the cells with SFM and incubated the cells with EDTA to dissociate all available cell surface annexin A2 and A5 ^7^. Cinnamycin treatment dramatically increased the amount of annexin A2 and A5 in the EDTA eluate without impacting cell morphology or viability, as shown by microscopy and lactate dehydrogenase activity in the eluate fraction, respectively (Fig. 1b and Supplementary Fig. 1a-b). As a negative control, no annexin A2 and A5 was detected in the eluate when cells were incubated in SFM without EDTA (Fig. 1b). The increase in annexin A2 and A5 on the cell surface in cinnamycin-treated cells was specific, as no cytosolic proteins or transmembrane proteins such as actin, Arf1, Arf6 and transferrin receptor were detected in the eluate fractions (Fig. 1b and Supplementary Fig. 1c). When cinnamycin was used at concentrations that compromised cell membrane integrity, we observed actin release in the eluate fraction (Supplementary Fig. 1d), suggesting that actin in the eluate correlated with cell lysis. Mass spectrometry confirmed that cinnamycin stimulated translocation of annexin A1, A2, A3, A4 and A5 to the cell surface (Fig. 1c). This phenomenon was not limited to HeLa cells and could be demonstrated in several other lines (Supplementary Fig. 1e). Furthermore, mastoparan X, another lipid-flipping toxin ^8^, caused a similar increase in annexin A2 detectable on the cell surface (Supplementary Fig. 1f).

**Figure 1.**
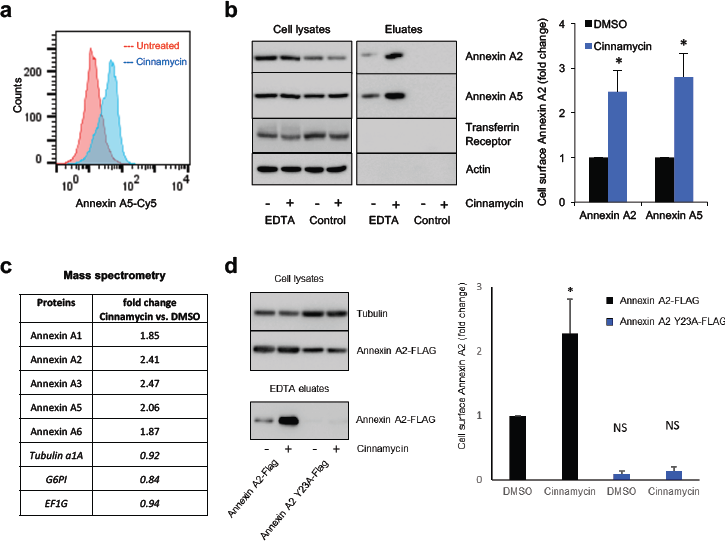
Cinnamycin facilitates annexin translocation across membranes in cells. **(a)** HeLa cells were treated with 1 µM cinnamycin for 50 min at 37°C. Then, annexin A5-Cy5 and propidium iodide (PI) were added and incubated for a further 10 min at 37°C. Annexin A5-Cy5 binding and PI accumulation were analysed by FACS. Representative histograms of annexin A5-Cy5 binding to live cells are shown (*n*=3). **(b)** Western blotting analysis of cell lysates and eluates of HeLa cells treated with cinnamycin (30 min at 37°C) and then with EDTA (10 min at 37°C) as indicated. Quantification of cell surface annexin A2 and annexin A5 (fold change measured as band intensity [cinnamycin(eluate/lysate)/DMSO(eluate/lysate)]) is shown. Error bars represent ±s.e.m. from biological replicates (*n* = 3); * *p*<0.05. **(c)** Mass spectrometry analysis of cell surface annexin from the samples in (E). Data are fold change of the number of peptides identified measured as [cinnamycin(eluate/lysate)/DMSO(eluate/lysate)]. **(d)** Left, western blotting analysis of cell lysates and eluates of HeLa cells transfected for 24 h with annexin A2-FLAG or annexin A2 Y23A-FLAG and then treated with cinnamycin (30 min at 37°C) and with EDTA (10 min at 37°C) as indicated. Right, quantification of cell surface annexin A2 (fold change measured as band intensity [cinnamycin(eluate/lysate)/DMSO(eluate/lysate)]). Error bars represent ±s.e.m. from biological replicates (*n* = 3); * *p*<0.05, NS: not significant.

To assess the requirement for lipid flipping activity in annexin translocation to the cell surface, we conducted the assay at 4°C. At this temperature, cinnamycin binds to the membrane but lipid flipping is abrogated ^5^. Cinnamycin-mediated annexin A2 translocation to the cell surface was inhibited at 4°C (Supplementary Fig. 1g). To evaluate the importance of annexin A2 lipid binding in the transport process, we analysed the translocation of an annexin A2 mutant (Y23A) which is defective in lipid binding and has a defect in cell surface localisation ^1^. Cinnamycin was unable to facilitate the translocation of the annexin A2 mutant (Fig. 1d). Together, these data support a mechanism whereby annexin A2 and A5 are transported to the cell surface by first binding to lipids on the inner leaflet of the membrane before being translocated across to the cell surface during lipid remodelling.

To evaluate whether lipid flipping is the minimal requirement for annexin transport across membranes, we developed an *in-vitro* liposome system using recombinant annexin A5 and cinnamycin. We first confirmed that cinnamycin flips lipids in liposomes using an assay based on the quenching of NBD-PE by dithionite (Supplementary Fig. 2). Next, liposome binding and sedimentation experiments showed that annexin A5 binds to phosphatidylcholine:phosphatidylethanolamine (PC:PE) liposomes in the presence of calcium and can be removed from the membrane by the calcium chelator EGTA (Fig. 2a, lane 1 vs 3). Interestingly, pre-treatment with cinnamycin increased the EGTA-resistant annexin A5 fraction (Fig. 2a, lane 4). This suggests that annexin A5 was translocated from the outer leaflet of the liposome membrane (surface of the liposome) to the inner leaflet or lumen of the liposome where it was protected from removal by EGTA. To confirm this phenomenon, we used a proteinase K protection assay. Proteinase K is very efficient in cleaving exposed proteins from membrane surfaces ^9^. PC:PE liposomes were pre-incubated with annexin A5 before incubation with proteinase K, which resulted in digestion of free and surface-bound annexin A5 detected by a dramatic decrease in amount of full-length annexin A5 seen by western blot (Fig. 2b, lane 1 vs 3). A small fraction of full-length annexin A5 and a near full-length cleavage product was also detected; this corresponded to fully-protected and partially membrane-inserted annexin A5, respectively (Fig. 2b, lane 3). Protease protection was membrane-dependent, as all available annexin A5 was degraded in the presence of membrane-solubilising detergent (triton) (Fig. 2b, lane 4). Importantly, cinnamycin pre-treatment increased the levels of the full-length annexin A5 and the near full-length cleavage product detectable after proteinase K digestion (Fig. 2b, lane 5). Therefore, cinnamycin increased the proportion of annexin A5 protected from proteinase K degradation, in agreement with the EGTA assay.

**Figure 2.**
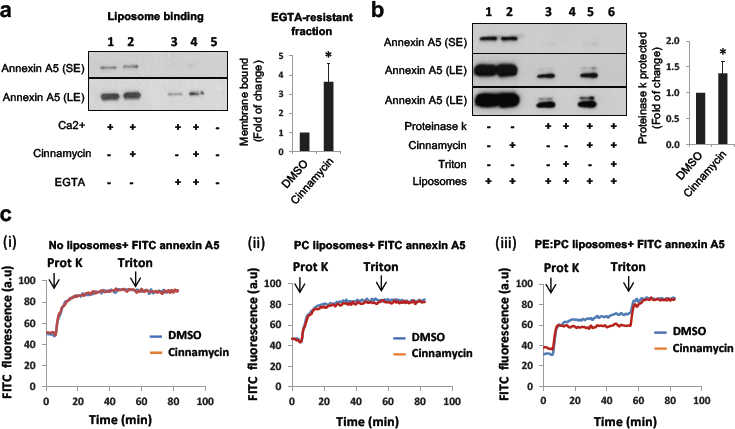
Cinnamycin facilitates annexin translocation across membranes in liposomes. **(a)** Cinnamycin increases EGTA-resistant membrane-bound fraction of Annexin A5. PC:PE liposomes (MLV) in CaCl_2_-containing buffer were incubated with Annexin A5. Cinnamycin or DMSO was added for a further 40 min incubation at 37°C. Some of the samples were also treated with EGTA before all samples were centrifuged. Liposome pellets were mixed with boiling SDS-sample buffer, separated by SDS-PAGE and analysed by western blot with anti-annexin A5 antibodies for the detection of the membrane-bound fraction of Annexin A5. Results are mean ± s.d. comparing lane 3 vs. lane 4, n=3 independent experiments, * *P <* 0.05 two-tailed paired t-test. (**b**) Protease protection assay (western blot). PC:PE liposomes (LUV) in CaCl_2_ containing buffer were incubated with FITC-Annexin A5. Cinnamycin or DMSO was added for further 40 min incubation at 37°C. Proteinase K alone or together with triton X-100 (0.5%) was added to liposome samples for further 1 hr at 37°C. After protease inactivation and boiling with SDS sample buffer, samples were resolved by SDS-PAGE and the proteinase K-protected fraction of Annexin A5 was detected by western blot analysis with anti-annexin A5 antibodies. Results are mean ± s.d. comparing lane 3 vs. lane 5, n=4 independent experiments, * *P <* 0.05 two-tailed paired t-test‥ SE= short exposure; LE = long exposure **(c)** Protease protection assay analysing proteinase K-induced FITC-annexin A5 dequenching. FITC-annexin A5 was added to buffer alone (i), PC alone LUV suspension (ii) and PC:PE LUV suspension (iii). Cinnamycin or DMSO was added to the samples and changes in FITC fluorescence during the experiment time were recorded. After 40 min, proteinase K was added to the samples for further 50 min at 37°C. Then, triton X-100 (0.5%) was added to the proteolysis samples for further 25 min. Results are average of duplicate measurements. Similar effects were detected in two independent experiments.

A similar protection assay was used in combination with a fluorescence quenching assay, allowing a real-time readout for degradation. In this assay, annexin A5 was labelled with fluorescein isothiocyanate (FITC) molecules. At high density, the fluorescence of the FITC molecules is quenched due to their close proximity on annexin A5. When proteinase K is added, accessible FITC-annexin A5 is cleaved, allowing the FITC molecules to be spatially separated, triggering an increase in their intrinsic fluorescence (dequenching). Proteinase K-induced FITC-annexin A5 dequenching was unaffected when incubated with cinnamycin then triton without liposomes, or when incubated with PC-only liposomes (Fig. 2c (i-ii)), as annexin A5 does not bind PC alone ^10^. By contrast, proteinase K-induced FITC-annexin A5 dequenching was reduced when PE-containing liposomes were used (Fig. 2c(iii)). This is consistent with results from the previous protection assay, indicating that there is a fraction of annexin A5 protected from protease cleavage (Fig. 2c(iii)). Furthermore, in agreement with previous results, PE-containing liposomes pre-treated with cinnamycin exhibited further attenuated FITC dequenching, compared to control/DMSO (Fig. 2c(iii)). Finally, the addition of triton completely recovered FITC fluorescence in both DMSO and cinnamycin conditions (Fig. 2c (iii)). Taken together, these three approaches suggest that lipid flipping induced by cinnamycin facilitates annexin A5 translocation across liposome membranes.

Given that cinnamycin is sufficient to translocate annexin across membranes in cells and liposomes, we looked for a mammalian protein that could drive this process in cells. Plasma membrane lipid asymmetry is maintained by transmembrane proteins that flip lipids from the inner leaflet to the outer leaflet and vice versa ^11,12,13^. One family of proteins with lipid flipping activity are the scramblases ^14^. Scramblase activity is calcium-dependent and energy-independent ^13-15^. Scramblase dysfunction results in Scott’s syndrome, a mild bleeding disorder resulting from a lack of phosphatidylserine externalisation ^16^. Scott’s syndrome has been attributed to mutations in the phospholipid scramblase known as TMEM16F (also called anotamin-6) ^15^. Due to the importance of TMEM16F lipid flipping activity *in vivo,* we investigated a role for TMEM16F in the translocation of annexin A2 and A5 to the cell surface. Clonal TMEM16F HeLa knockout cell lines were generated using CRISPR/Cas9 and matched wild-type controls with no targeting were also isolated (Supplementary Fig. 3a, b). We confirmed gene targeting using sequencing and qPCR (Supplementary Fig. 3c). We were unable to measure the protein level of TMEM16F as the few antibodies that we tried did not show specific signals on western blot (data not shown). However, we confirmed a functional defect in lipid flipping in the TMEM16F knockout cell lines by measuring the level of PS on cell surface in cells challenged with ionomycin (Fig. 3a(i)). We treated TMEM16F wild-type or -deficient cells with ionomycin for 10 min at 37°C in the presence of annexin A5-Cy5 and propidium iodide. Cell surface PS (and PE) were analysed by measuring the amount of annexin A5-Cy5 bound to live cells by flow cytometry. TMEM16F-deficient cells were unable to externalise PS in response to an increase in intracellular calcium, whereas wild-type cells and positive matched non-targeted controls showed an increase in the amount of PS on the cell surface (Fig. 3a(i) and Supplementary Fig. 4a). This confirms that TMEM16F activity is abolished in TMEM16F knockout cells. Interestingly, the level of PS (and PE) on the cell surface of TMEM16F deficient cells under unstimulated conditions was also slightly reduced compared to the wild-type and untargeted controls (Fig. 3a(ii) and Supplementary Fig. 4b), showing for the first time that TMEM16F is constitutively active.

**Figure 3.**
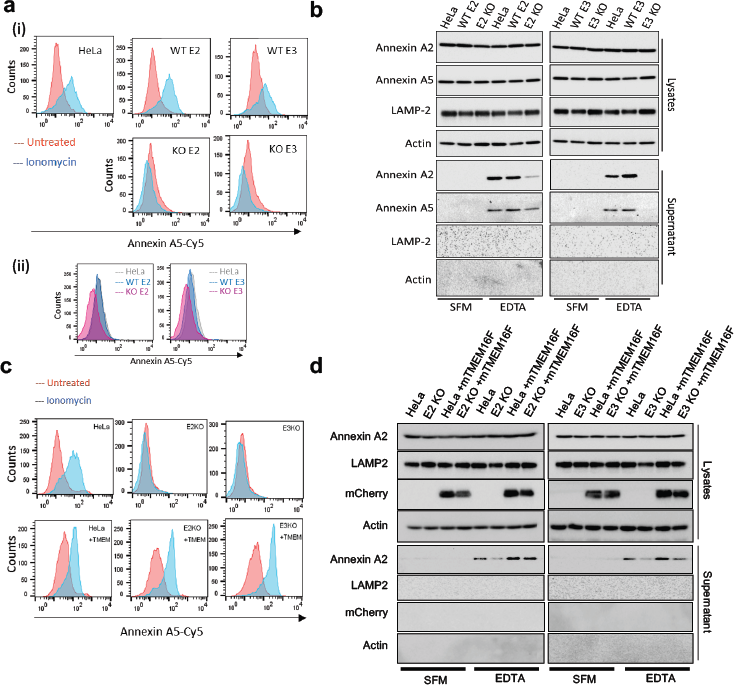
TMEM16F lipid flipping activity is required for annexin A2 and A5 cell surface localisation. **(a)** (i) Wild type, positive control and TMEM16F knockout cells were treated with 10 mM ionomycin for 10 min at 37°C in the presence of annexin A5-Cy5 and PI. Annexin A5-Cy5 binding and PI accumulation were analysed by FACS. Representative histograms of annexin A5-Cy5 binding to live cells are shown (*n*=4). (i) Basal cell surface PS is reduced in TMEM16F knockout cells. Cells were incubated with annexin A5-Cy5 and PI for 10 min at 37°C before FACS analysis. Representative histograms displayed (*n*=5). **(b)** Wild-type, positive control and TMEM knockout HeLa cells were incubated in versene (EDTA solution) for 10 min at 37°C before the eluate collected and analysed for annexin A2 and A5 by western blotting. Representative western blot shown (*n*=4). **(c)** Wild-type and TMEM16F-knockout Hela cells alone or expressing mCherry-mTMEM16F were treated with ionomycin and analysed for annexin A5-cy5 binding as described in (a). Representative experiment shown (*n*=3). **(d)** Annexin A2 and A5 on the cell surface were evaluated with EDTA and western blotting as described in (C). Representative experiment shown (*n*=4).

To investigate the role of TMEM16F in translocation of annexins, we assessed the level of annexin A2 and A5 on the cell surface of TMEM16F deficient cells. Cells were treated with EDTA to release annexin A2 and A5 and the eluate was assessed by western blot, as described earlier. Strikingly, in both TMEM16F-deficient clones, annexin A2 and A5 were severely reduced in the EDTA eluate (Fig. 3b). Annexin A2 and A5 were not detected in the SFM eluate, and actin and LAMP-2 were absent from all eluates (SFM or EDTA) indicating this was a specific process (Fig. 3b). To ensure that the lack of annexin on the cell surface was due to reduced translocation rather than reduced retention at the cell surface, we assessed the level of annexin released into the supernatant over 24 hours. No annexin A2 and A5 were detected in the supernatant from wild-type or TMEM16F-deficient cells (Supplementary Fig. 4c). This demonstrated that annexin A2 and A5 were not translocated and thus present in the medium in TMEM16F-deficient cells (Supplementary Fig. 4c), whereas annexin A2 and A5 are translocated across the membrane in wild-type cells and detectable on the cell surface (Fig. 3b). The lack of annexin A2 and A5 on the cell surface in TMEM16F-deficient cells is not due to off-target effects as the phenotype is consistent across both clones targeted with different sgRNAs. Furthermore, the phenotype was normalised when the TMEM16F-deficient cells were reconstituted with mouse, mCherry-tagged TMEM16F (mCherry-mTMEM16F) that restored lipid flipping activity (Fig. 3c, d and Supplementary Fig. 5a, b and c). This established that TMEM16F is required for transport of annexin to the cell surface.

Our data indicate that cinnamycin stimulates lipid flipping in a manner comparable to TMEM16F, therefore, we set out to determine if its activity could substitute for TMEM16F. Cells were treated with cinnamycin and the level of PS and PE externalisation was assessed by annexin A5-Cy5 binding to live cells by flow cytometry. Cinnamycin stimulated lipid flipping in both wild-type and TMEM16F-deficient cells (Fig. 4a and Supplementary Fig. 6a, b). Therefore, cinnamycin can be used as a surrogate for TMEM16F lipid flipping activity. TMEM16F-deficient cells treated with DMSO showed reduced annexin A2 and A5 in the EDTA eluate (Fig. 4b), however when treated with cinnamycin there was a significant increase in the amounts of annexin A2 and A5 detectable in the EDTA eluate in these cells (Fig. 4b). Cinnamycin also increased the level of annexin A2 and A5 on the cell surface in wild-type HeLa, as expected (Fig. 4b). This demonstrates that lipid flipping activity is sufficient for translocation of annexin A2 and A5 from the cytosol to the cell surface and was not due to a lack of annexin retention at the cell surface as annexin A2 was not present in the medium during cinnamycin treatment as described earlier (Supplementary Fig. 6c).

**Figure 4.**
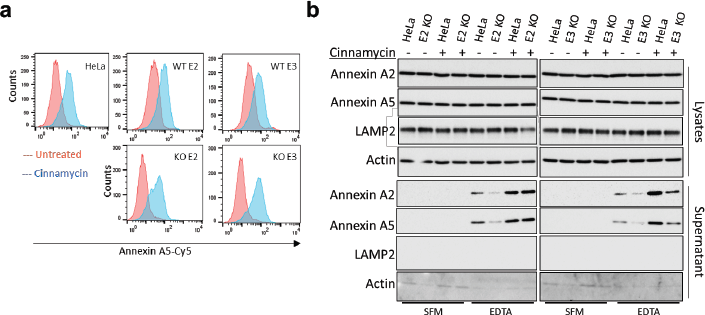
Cinnamycin restores annexin A2 and A5 cell surface localisation in TMEM16F-deficient cells. **(a)** Wild type, positive control and TMEM16F-knockout cells were treated with 1 µM cinnamycin for 50 min at 37°C, annexin A5-Cy5 and PI were then added and incubated for a further 10 min at 37°C. Annexin A5-Cy5 binding and PI accumulation were analysed by FACS. Representative histograms of annexin A5-Cy5 binding to live cells are shown (*n*=3). **(b)** Wildtype, positive control and TMEM-knockout HeLa cells were incubated with DMSO or cinnamycin at 37°C for 1 hour. Cells were washed and annexin A2 and A5 released with versene (EDTA solution) for 10 min at 37°C before the eluate collected and analysed by western blotting. Representative western blot shown (*n*=4).

In summary, we have identified a mechanism for membrane translocation of annexins which is mediated by lipid flipping/remodelling. This process provides an explanation for the longstanding mystery of how some cytoplasmic proteins reach the cell surface independent of conventional secretion or described modes of unconventional secretion ^17-20^. In cells, this lipid flipping-dependent translocation is primarily mediated by TMEM16F and enables transport of a protein from the inner intracytoplasmic domain of the plasma membrane to the exterior outer leaflet. This transport is ATP-independent and calcium-dependent and does not require the complex translocon-type machinery as seen for conventional trafficking of proteins into the ER or mitochondria.

## Methods

### Cell culture

HeLa cells (from ATCC) were cultured in DMEM D6546 (Molecular Probes) containing 10% fetal bovine serum, supplemented with 2 mM L-glutamine and 100 U ml–1 penicillin/streptomycin in 5% CO_2_ at 37°C. PANC-1 and AsPC-1 (a gift from Frances Richards, Cancer Research UK Cambridge Institute) were cultured in DMEM D6546 (Molecular Probes) containing 10% fetal bovine serum, supplemented with 2 mM L-glutamine and 100 U ml–1 penicillin/streptomycin in 5% CO_2_ at 37°C, or RPMI-1640 (Molecular Probes) containing 10% fetal bovine serum, supplemented with 2 mM L-glutamine and 100 U ml–1 penicillin/streptomycin in 5% CO2 at 37°C, respectively. Human embryonic kidney cells 293 (HEK) were cultured in DMEM D6546 (Molecular Probes) containing 10% fetal bovine serum, supplemented with 2 mM L-glutamine and 100 U ml–1 penicillin/streptomycin in 5% CO_2_ at 37°C. All cells were tested negative for mycoplasma contamination.

### Antibodies

Antibodies include: mouse monoclonal anti-annexin A2 (BD Biosciences; 610071; 1/1,000), mouse monoclonal anti-annexin A5 (Abcam; ab54775), mouse monoclonal anti-transferrin receptor (Zymed; H68.4; 1/1,000), rabbit polyclonal anti-actin (Sigma; A2066; 1/2,000), mouse monoclonal anti-tubulin (Sigma; T9026; 1/4,000), mouse monoclonal anti-FLAG (Sigma-Aldrich; clone M2; 1/4,000), mouse monoclonal anti-Arf1 (Santa Cruz Biotechnology; sc-53168), mouse monoclonal anti-Arf6 (Santa Cruz Biotechnology; clone 3A-1; 1/500), mouse monoclonal anti-LAMP2 (Biolegend; 354302; WB: 1/1,000) and rabbit polyclonal anti-mCherry antibody (Genetex; GTX128508-S).

### Reagents

Reagents include: cinnamycin (Santa Cruz Biotechnology; sc-391464), mastoparan X (Alfa Aesar; J61173), annexin A5-FITC (Abcam; ab14085), 1,2-Dioleoyl-sn-glycero-3-phosphoethanolamine (DOPE, Sigma-Aldrich; 54008), 2-Oleoyl-1-palmitoyl-sn-glycero-3-phosphocholine (POPC, Sigma-Aldrich; P3017), 1,2-dioleoyl-sn-glycero-3-phosphoethanolamine-N-(7-nitro-2-1,3-benzoxadiazol-4-yl) (NBD-PE, Avanti Polar Lipids, 810145) and proteinase K (Molecular Biology; BP1700-100), EGTA (Sigma-Aldrich; E3889), Sodium dithionite (Sigma-Aldrich; 71699), versene solution containing ethylenediaminetetraacetic acid (EDTA) (Gibco; 15040-033), propidium iodide solution (Biolegend; 421301), QuickExtract DNA extraction solution (Epicenter; QE0905T), Herculase II fusion DNA polymerase (Agilent; 600675), annexin V-FITC and annexin V-Cy5 Apoptosis Staining / Detection Kit (ab14085, ab14150) and ionomycin (Cayman Chemical Company; 10004974). Oligonucleotides for TMEM16F CRISPR targeting and sequencing were synthesized from Sigma-Aldrich (Supplementary Table 1).

### Plasmids

Annexin A2-FLAG and Annexin A2 Y23A-FLAG were a gift from Lei Zheng (Johns Hopkins Technology Ventures) ^1^, pSpCas9(BB)-2A-Puro (PX459) was a gift from Feng Zhang (Addgene plasmid # 48139). ANO6-Plvx-mCherry-c1 was a gift from Renzhi Han (Addgene plasmid # 62554). psPAX2 and pMD2.G were a gift from Didier Trono (Addgene plasmid # 12260 and # 12259, respectively).

### Preparation of liposomes

Thin films were generated following dissolution of the lipids in a 2:1 (v/v) chloroform/methanol mixture and then dried under a stream of argon gas while they were rotated. The final compositions in mole percentage of the different liposome population were PE-containing liposomes (50% DOPE and 50% POPC) and PC-containing liposomes (100% POPC). The films were lyophilized overnight, and the containers were sealed with argon gas to prevent oxidation and stored at −20°C. Multilamellar vesicles (MLV) were generated by solubilising the lipid films with physiologic salt buffer (PSB), composed of 100 mM KOAc, 2 mM Mg(OAc)_2_, 50 mM HEPES, pH 7.4, using vigorous vortexing. For generation of large unilamellar vesicle (LUV), the films were suspended in PSB and vortexed for 1.5 min. The lipid suspension underwent five cycles of freezing and thawing followed by extrusion through polycarbonate membranes with 1 and 0.1 µm diameter pores (from Avanti Polar Lipids) to create LUV.

### Lipid flipping assay in liposomes

The following lipid flipping assay in liposomes is well established, see ref ^21^. Briefly, fluorescent measurements were performed using SpectraMax M5 (Molecular Devices) in 96-well plates with total reaction volumes of 100 µl at a constant temperature of 37°C. Excitation was set on 480nm, and emission was set on 530 nm (NBD fluorescence) with low PMT sensitivity. LUV (PC:PE 1:1, and 0.6% NBD-PE) at 500 µM in PSB were pre-incubated with cinnamycin (10 µM) or DMSO, as a control, for 30 min and changes in NBD fluorescence during the experimental period were recorded and the effects of dithionite (3 mM) were assessed. Dithionite reduces the NBD molecules on the head group of the lipids. Because only NBD in the outer membrane leaflet is accessible to react with dithionite, a partial decrease in fluorescence is observed. When the fluorescent system reached a steady-state, a membrane-solubilising detergent (triton 0.5%) was added, which exposes the NBD in the inner leaflet and the decrease in NBD fluorescence was assessed. If cinnamycin induces lipid flipping, then there would be a difference in the level of fluorescence (compared to DMSO control) after dithionite treatment, which correlates to the different level of NBD-PE in the outer leaflet.

### Annexin A5 liposome binding experiments

MLV suspension (1 mM) was supplemented with CaCl2 (5 mM) and was incubated with FITC-Annexin A5 (1:7 dilution from stock solution) for 30 min in room temperature to allow Annexin A5 binding to liposomes. Cinnamycin (10 µM) or DMSO was added for a further 40 min incubation at 37°C. Some of the samples were also treated with EGTA (10 mM) for 20 min in in room temperature before all samples were centrifuged (16,000g, 30 min at 4°C). Liposome pellets were mixed with boiling SDS-sample buffer for 1 min, separated by SDS-PAGE, transferred onto PVDF membranes and subjected to western blot analysis with anti-Annexin A5 antibodies for the detection of the membrane-bound fraction of Annexin A5.

### Protease protection assay - SDS-PAGE analysis

LUV suspension (100 µM) was supplemented with CaCl_2_ (5 mM) and was incubated with FITC-Annexin A5 (1:4.5 dilution from stock solution) for 30 min in room temperature. Cinnamycin (10 µM) or DMSO was added for further 40 min incubation at 37°C. For proteinase K experiments, a stock solution of the protease was freshly prepared by in PSB (10 mg/ml) prior to the experiment and was kept on ice. Proteinase K was added to liposome samples (1:20 dilution from stock solution) and incubation was carried out for further 1 h at 37°C to allow full digestion. For some samples, triton X-100 (0.5%) was added together with Proteinase K. We then inactivated the protease by adding a small amount of PMSF (from 0.25 M stock dissolved in DMSO) to the liposome samples, and put this on ice. The proteolysis samples were transferred into boiling SDS sample buffer and immediately pipetted several times. The boiled protein samples were resolved by SDS-PAGE and the proteinase K-protected fraction of Annexin A5 was detected by western blot analysis with anti-Annexin A5 antibodies.

### Protease protection assay – fluorescent measurements

Fluorescent measurements were performed using SpectraMax M5 (Molecular Devices) in 96-well plates with total reaction volume of 100 µl in constant temperature of 37°C. All measurements were performed in duplicate. Excitation was set on 480nm, and emission was set on 530 nm (detection of FITC fluorescence) with low PMT sensitivity. FITC-Annexin A5 (1:25 dilution from stock solution) was added to LUV suspension (500 µM) supplemented with CaCl_2_ (5 mM) and divided in 100 µl samples in a 96-well plate. Cinnamycin (10 µM) or DMSO was added to the plate and FITC fluorescence was recorded every min using the kinetic measurement mode of the instrument. After 40 min, proteinase k was added to the wells for further 50 min incubation and FITC florescence was recorded. Then, triton X-100 (0.5%) was added to the proteolysis samples for further 25 min.

### Cinnamycin treatment and EDTA assay in cells

Cells were treated from 30 min to 1 h at 37°C with 1 µm cinnamycin (diluted in serum-free medium, SFM) or DMSO used as control. Cells were then washed twice with SFM and incubated with versene (EDTA solution) or SFM used as control for 10 min at 37°C. The supernatant was collected, floating cells pelleted at 300 *g* for 5 min before filtering through a 0.2 µm syringe filter. The a sample of clarified supernatant was then mixed with 4X sample buffer (50 mM Tris-HCl pH 6.8, 2% SDS (w/v), 0.1% Bromophenol Blue, 10% Glycerol and 100 mM DTT) and boiled for 5 min. Cells were lysed in lysis buffer (20 mM Tris-HCl pH 6.8, 137 mM NaCl, 1 mM EDTA, 1% triton X-100 and 10% glycerol) at 4°C for 10 min and insoluble material removed by centrifugation at 10,000 *g* for 10 min 4°C. Sample buffer was added and cell lysate were samples boiled added (as above). Cell lysates and cell supernatants were then subjected to SDS-PAGE.

### Western blotting

All samples were resolved by 12% SDS-polyacrylamide gel electrophoresis (PAGE) and transferred to polyvinylidene difluoride membranes for blotting. Membranes were blocked with 0.05% (w/v) skim milk powder in PBS containing 0.1% Tween-20 (PBS-Tween) for 30 min at room temperature. Membranes were then probed with an appropriate dilution of primary antibody overnight at 4°C. Membranes were washed three times in PBS-Tween before incubation in diluted secondary antibody for 1 h at room temperature. Membranes were washed as before and developed with ECL (Amersham ECL Western Blotting Detection Reagent RPN2106 for the detection of proteins in the cell lysates or Cyanagen, Westar XLS100 for the detection of proteins in the eluate fractions) using a Bio Rad Chemi Doc XRS system. Membranes were stripped with Restore plus (ThermoFisher Scientific, 46430) as per manufactures instructions.

### Lentiviral transfection

HEK293FT packaging cells growing in 10-cm dishes were transfected with a mix of 11.68 µg packaging vector (psPAX2), 5.84 µg envelope vector (pMD2.G) and 18.25 µg ANO6-Plvx-mCherry-c1 vector. PEI (polyethylenimine) was used as transfection reagent. 48h after transfection, cell culture medium was collected and replaced by fresh medium; this step was repeated 2 times at intervals of 24 h. Virus preparations were then combined. Viral particles were added to cells, spin at 1,000g for 30 min and incubated overnight. After 24 h, medium was replaced by full medium and cells were incubated for 5 more days. Transduced cells were selected with puromycin and sorted to enrich mCherry-expressing cells.

### LDH assay

Lactate dehydrogenase (LDH) assay was performed according to manufacturer instructions (Thermo Fischer, 88953).

### Mass spectrometry

Samples were submitted to the Cambridge Institute for Medical Research–Institute of Metabolic Science proteomics facility where they were analyzed using Thermo Orbitrap Q Exactive with EASY-spray source and Dionex RSLC 3000 UPLC.

### TMEM16F CRISPR-mediated gene disruption

TMEM16F was targeted in either exon 2 or exon 3, both of which are conserved across splice variants. TMEM16F specific oligonucleotides (Sigma-Aldrich; table S1) were designed and top and bottom strands were annealed, and then cloned into the Cas9 expression vector pSpCas9(BB)-2A-Puro (PX459) (Addgene plasmid # 48139) as previously described ^22^. Transfected cells were selected with 2.5 µg/ml puromycin for 24 hours. Once recovered, cells were single cell sorted into 96 well plates by FACS. TMEM16F targeting was verified by collecting genomic DNA from clonal lines using QuickExtract DNA extraction solution and the CRISPR/Cas9 targeted region amplified with primers flanking at least 200 base pairs either side of the expected cut site (table S1). PCR products were sequenced by Sanger sequencing and insertions and deletions analysed by using the Tracking of Indels by DEcomposition (TIDE) web tool ^23^. Additionally, to analyse insertions and deletions larger than 50 base pairs, the R code was kindly provided by Prof van Steensel. TIDE analysis showed that the expected region had been targeted and each knockout clone was devoid of wild type TMEM16F DNA (Fig. S3A and B). Clonal wild type control lines that had been through transfection, selection and single cell cloning steps, but had not efficiently targeted TMEM16F, were used as matched positive controls for each exon (Fig. S3A and B). Due to the lack of antibodies specific for TMEM16F we were unable to analyse expression at the protein level, therefore, we assessed the mRNA levels which were reduced in TMEM16F knockout cells (Fig. S3C).

### Ionomycin and cinnamycin PS flow cytometry assay

Approximately 1×10^6^ HeLa cells in 6 well plates were washed once in serum-free medium, incubated in versene (EDTA solution) at 37°C until detached and collected in to an excess volume of complete DMEM. Cells were pelleted at 300 g and resuspended in 500 µl annexin V binding buffer (Abcam). Cells were then transferred into FACS tubes containing 5 µl annexin V-Cy5 and 1 µl propidium iodide and ether 1 µl ionomycin or ethanol. Cells were carefully mixed and incubated at 37°C for 10 min only. Cells were immediately analysed on a FACSCalibur (BD) equipped with lasers providing 488nm and 633nm excitation sources. Annexin-Cy5 fluorescence was detected in FL4 detector (661/16 BP) and propidium iodide was detected in FL2 (585/42 BP). For analysis of cinnamycin, cells were collected into annexin V binding buffer as above and incubated with 1 µM cinnamycin or the equivalent volume of DMSO at 37°C for 50 min. Annexin V-Cy5 and propidium iodide were then added and cells incubated at 37°C for a further 10 min. Cells were analysed by FACS immediately, as described above.

### Cell sorting

For sorting, cells were collected with trypsin/EDTA, washed and FACS was carried on an Influx cell sorter (BD) or Aria-Fusions (BD) equipped with lasers providing 488 nm and 640 nm excitation sources. mCherry fluorescence was detected in 610/20 BP detector on Influx and the Aria Fusion.

### Statistical analysis

Significance levels for comparisons between groups were determined with *t*- tests.

## Acknowledgements

We thank Frances Richards (Cancer Research UK Cambridge Institute) for the PANC1 and AsPC1 cells, Lei Zheng (Johns Hopkins Technology Ventures) for the Annexin A2 vectors. We also thank CIMR/IMS Proteomics Facility for the mass spectrometry analysis and the NIHR Cambridge BRC Cell Phenotyping Hub for cell sorting.

## Author contributions

AA, KM, and DCR designed liposome experiments, SES and KM designed cell line experiments. AA conducted liposome experiments. SES and KM conducted cell line experiments. AW conducted some of the experiments in cell. SES generated, validated and conducted TMEM16F experiments with support from KM. SES, AA, KM and DCR wrote the manuscript. KM and DCR supervised the research.

